# Long non-coding RNAs are largely dispensable for zebrafish embryogenesis, viability and fertility

**DOI:** 10.1101/374702

**Authors:** Mehdi Goudarzi, Kathryn Berg, Lindsey M. Pieper, Alexander F. Schier

## Abstract

Hundreds of long non-coding RNAs (lncRNAs) have been identified as potential regulators of gene expression, but their functions remain largely unknown. To study the role of lncRNAs during vertebrate development, we selected 25 zebrafish lncRNAs based on their conservation, expression profile or proximity to developmental regulators, and used CRISPR-Cas9 to generate 32 deletion alleles. We observed altered transcription of neighboring genes in some mutants, but none of the lncRNAs were required for embryogenesis, viability or fertility. Even RNAs with previously proposed non-coding functions (*cyrano* and *squint*) and other conserved lncRNAs (*gas5* and *lnc-setd1ba)* were dispensable. In one case (*lnc-phox2bb*), absence of putative DNA regulatory-elements, but not of the lncRNA transcript itself, resulted in abnormal development. LncRNAs might have redundant, subtle, or context-dependent roles, but extrapolation from our results suggests that the majority of individual zebrafish lncRNAs are dispensable for embryogenesis, viability and fertility.

---

Long non-coding RNAs (lncRNAs) comprise a heterogeneous group of transcripts longer than 200 nucleotides that do not encode proteins. LncRNAs have been proposed to affect the expression of neighboring or distant genes by acting as signaling, guiding, sequestering or scaffolding molecules^1–5^. The functions of specific lcnRNAs in dosage compensation (*xist*^6,7^, *tsix*^8^, *jpx*^9^) and imprinting (*Airn*^10,11^, *MEG3*^12,13^, *H19*^14,15^) are well established, and mutant studies in mouse have suggested that *fendrr, peril, mdget, linc-brn1b, linc-pint*^16^, and *upperhand*^17^ are essential for normal development. However, other studies have questioned the developmental relevance of several mouse lncRNAs, including *Hotair*^18^, *MIAT/Gumafu*^19^, *Evx1-as*^20^, *upperhand*, *braveheart* and *haunt*^21^. In zebrafish, morpholinos targeting the evolutionarily-conserved lncRNAs *megamind* (TUNA^22^) and *cyrano* resulted in embryonic defects^23^. However, a mutant study found no function for *megamind* and revealed that a *megamind* morpholino induced non-specific defects^24^. These conflicting results have led to a controversy about the importance of lncRNAs for vertebrate development^16,21^. We therefore decided to mutate a group of selected zebrafish lncRNAs using CRISPR-Cas9, and assay their roles in embryogenesis, viability and fertility.

Transcriptomic studies of early embryonic development^23,25^ and five adult tissues^26^ have identified over 2,000 lncRNAs in zebrafish^27^, of which 727 have been confirmed as non-coding based on ribosome occupancy patterns^28^. For our knockout study we selected 24 bona fide lncRNAs based on conservation, expression dynamics and proximity to developmental regulatory genes (see Fig 1 for selection criteria for each lncRNA). These criteria were chosen to increase the likelihood of functional requirement. In addition, we selected a protein-coding RNA with a proposed non-coding function (*squint*). The genomic location, neighbor-relationship, and expression levels of the selected lncRNAs and their neighboring genes are shown in supplementary figures Fig S1, Fig S2, and Fig S3, respectively.

**Figure 1:**
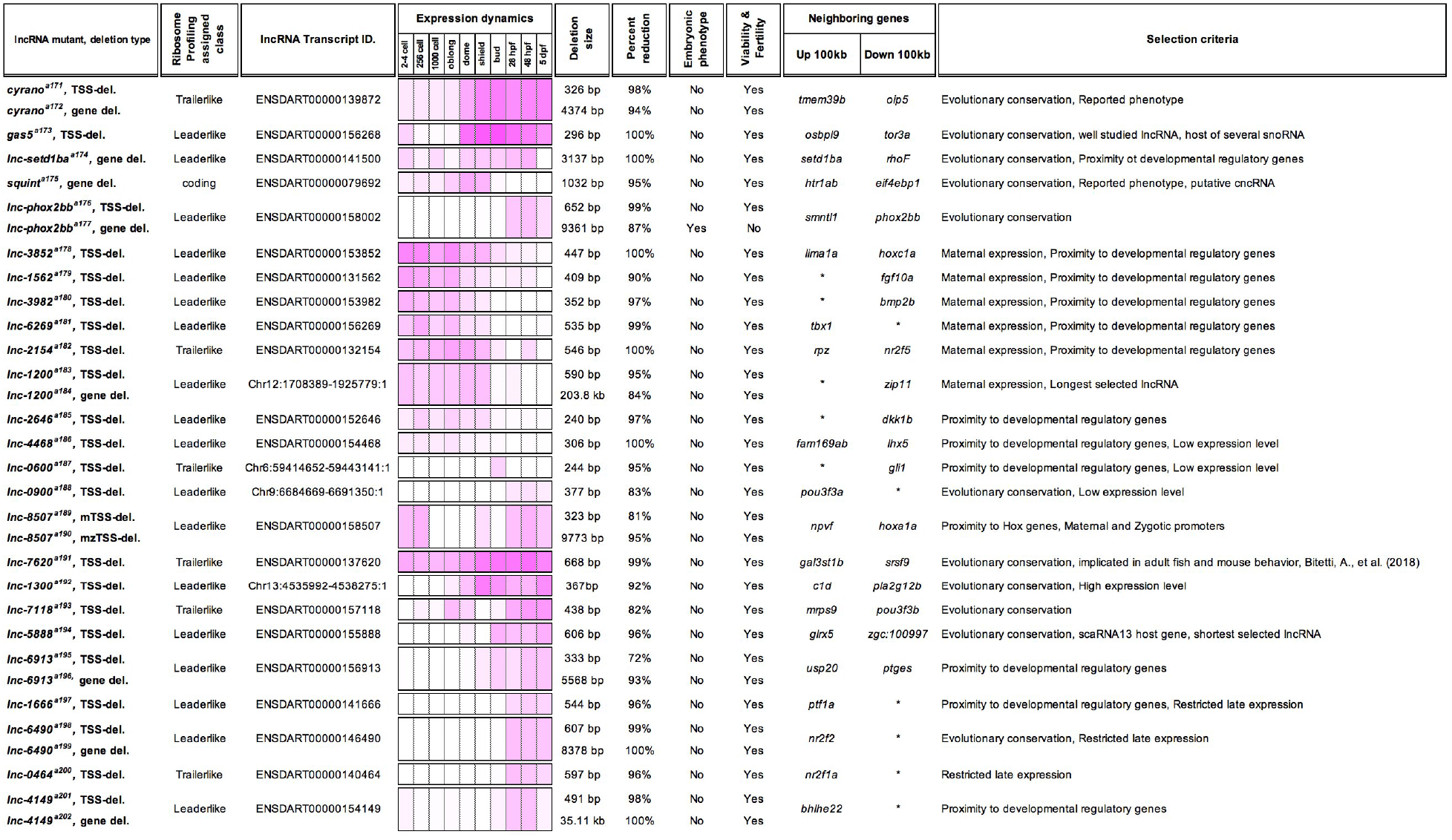
Summary of the lncRNA mutant study. lncRNA names are shown in the first column. lncRNAs were named using the last 4 digits of their corresponding ENSEMBL Transcript ID or their chromosome number if no transcript ID was available (e.g. lnc-1200 is located on chromosome 12). The second column represents ribosomal occupancy pattern along the length of lncRNAs in comparison to the 5’UTR, coding and 3’UTR of typical protein-coding transcripts^28^. The third column shows the transcript ID for the investigated lncRNA or its genomic coordinate in GRCz10. Column Four represents the expression dynamics of the lncRNAs (see also Figure S3) (log2 (FPKM +1) between 0 and 8)^25^. The fifth column shows the deletion size. Sixth column represent the percentage decrease in the level of lncRNA in comparison to wild type from 3 biological replicates (qRT-PCR). The seven and eight columns show the presence of embryonic phenotypes, viability and fertility (at least 15 adult pairs per allele) of homozygous mutant fish. Ninth column shows the upstream and downstream neighboring genes in a 200kb window centered around the lncRNA’s TSS. The last column provides the selection criteria for each lncRNA.

Using CRISPR-Cas9 (Fig S4) we generated 32 knockout-alleles. 24 alleles removed regions containing transcription start sites (TSS-deletion; 244bp to 736bp), and 8 alleles fully or partially removed the gene (1kb to 203kb) (Fig 1). qRT-PCR analysis demonstrated effective reduction in the levels of the targeted lncRNA transcripts (average reduction of 94 ± 6%; Fig 1), which was further tested and confirmed for a subset of lncRNAs by in situ RNA hybridization (Fig 2B, 3B, 4C, 4D, 5B and 6D).

**Figure 2:**
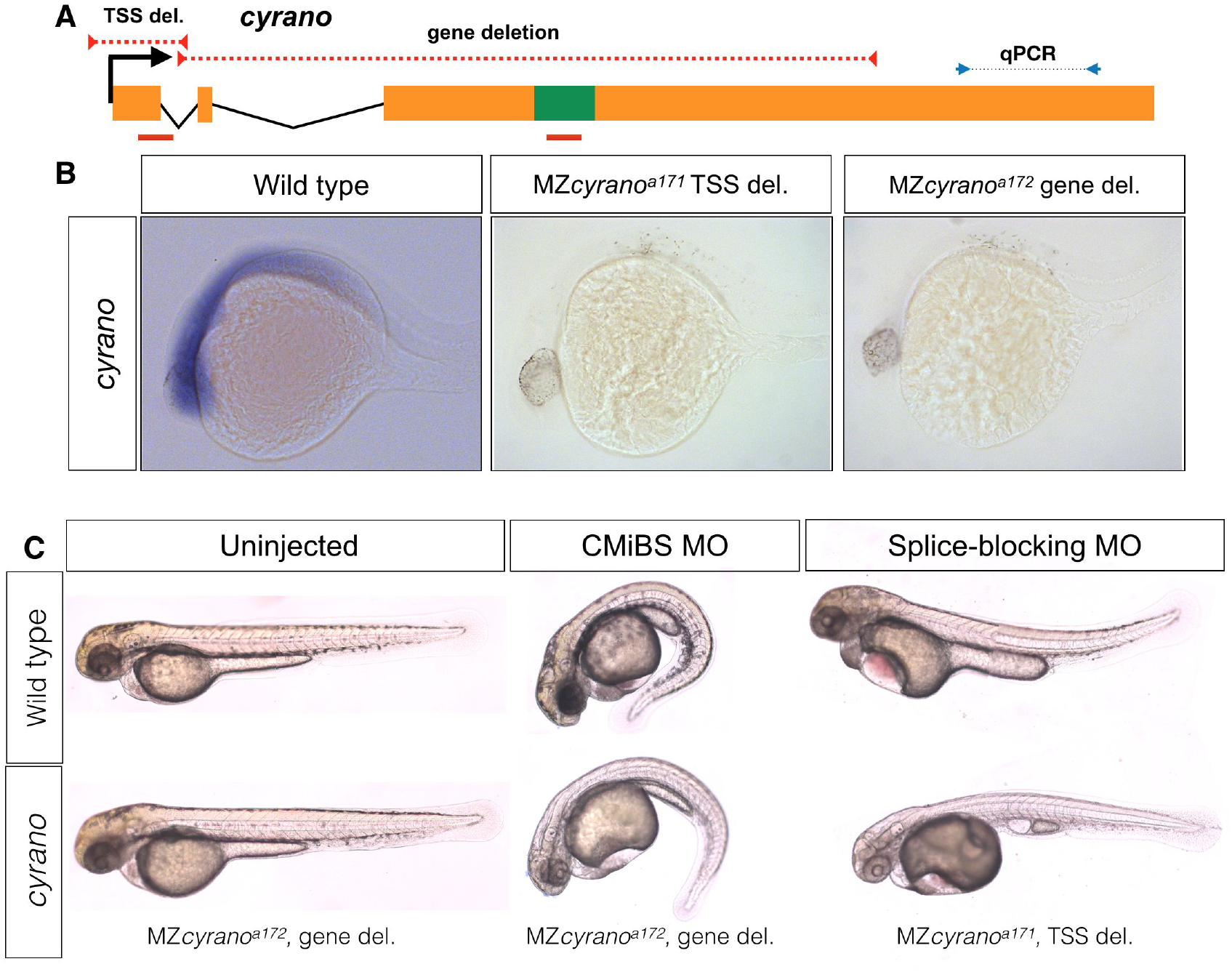
Normal embryogenesis of *cyrano* mutants. A) The positions of TSS-deletion allele and gene deletion allele are marked by dashed red lines. Green box represents the conserved element in *cyrano* which is complementary to *miR-7*. Solid red lines indicate the position of the first exon-intron boundary (e1i1) morpholino and conserved microRNA binding site (CMiBS) morpholinos. Arrows flanking black dotted line mark the primer binging sites for qRT-PCR product. B) Representative images of in situ hybridization for *cyrano* in wild type (15/15) and both homozygous TSS-deletion (21/22) and gene deletion (18/18). C) At 1-dpf gene deletion mutants (lower-left), (and TSS-deletion mutants, not shown) were not different from the wild type embryos (upper-left). Morpholino injected wild-type embryos (upper-middle and upper-left) reproduced observed phenotype in Ulitsky et. al.^24^. Morpholino injected deletion-mutants, lacking the corresponding binding sites for morpholinos, (lower-middle and lower-left) were comparable to morpholino injected wild types.

Previous observations in mammalian cell culture systems suggested that lncRNA promoters can affect the expression of nearby genes^29^. To test if these results hold true in vivo, we measured the changes in the expression of neighboring genes (a 200 kb window centered on each lncRNA) in lncRNA mutants. Several mutants displayed changes in the expression of neighboring genes (Fig S5). In particular, 10 out of 40 neighboring genes showed more than two-fold changes in expression, lending in vivo support to observations in cell culture systems^29^.

To determine the developmental roles of our selected lncRNAs, we generated maternal-zygotic mutant embryos (lacking both maternal and zygotic lncRNA activity) and analyzed morphology from gastrulation to larval stages, when all major organs have formed. Previous large-scale screens^30,31^ have shown that the visual assessment of live embryos and larvae is a powerful and efficient approach to identify mutant phenotypes, ranging from gastrulation movements and axis formation to the formation of brain, spinal cord, floor plate, notochord, somites, eyes, ears, heart, blood, pigmentation, vessels, kidney, pharyngeal arches, head skeleton, liver, and gut. No notable abnormalities were detected in 31/32 mutants. Moreover, these 31 mutants survived to adulthood, indicating functional organ physiology, and were fertile (Fig1). In the following section we describe the results for five specific lncRNAs and put them in the context of previous studies.

## Cyrano

*cyrano* is evolutionarily conserved lncRNA and based on morpholino studies, has been suggested to have essential functions during zebrafish embryogenesis^23^ and brain morphogenesis ^32^. c*yrano* has also been suggested to act as a sponge (decoy-factor) for HuR during neuronal proliferation^33^, regulate *miR-7* mediated embryonic stem cell differentiation^34^, and control the level of *miR-7* in the adult mouse brain^35^. We generated two mutant alleles that removed the TSS (*cyrano*^a171^) or the gene (*cyrano*^a172^), including the highly conserved *miR-7* binding-site (Fig 2A, B). The expression level of the nearby gene (*oip5*) was not affected in either of these mutants (Fig S5). In contrast to previous morpholino studies in zebrafish^23^ but in support of recent findings in mouse^35^, *cyrano* mutants developed normally and were viable and fertile.

The difference between morphant^23^ and mutant phenotypes might be caused by compensation in the mutants^36,37^. To test this possibility, we injected the previously used morpholinos targeting the first exon-intron boundary (e1i1) or the conserved *miR-7* binding site (CMiBS) into wild type and homozygous deletion mutants. The TSS-mutant allele lacked the e1i1 morpholino binding site and the gene deletion allele lacked the CMiBS morpholino binding site (Fig 2A). The previously reported phenotypes, including small heads and eyes, heart edema, and kinked tails were found in both wild type and mutants (Fig 2C), demonstrating that the morpholino-induced phenotypes were non-specific. These results reveal that *cyrano* transcripts or their evolutionarily conserved *miR-7* binding site, are not required for embryogenesis, viability or fertility.

## gas5

*gas5* is an evolutionarily conserved lncRNA *(growth-arrest specific 5)*^38^ that is highly expressed in early development (Fig 3B) and hosts several snoRNAs implicated in zebrafish development^39^. Knockdown and knockout studies in cell culture^40^ have indicated that *gas5* might act as a tumor suppressor^41^ and exert effects at distant genomic sites^42^. However, the role of this lncRNA in development has not been studied in any vertebrate. Our *gas^5a173^* mutant allele removed the sequences containing the TSS (−169 to +127) (Fig 3A) and resulted in complete elimination of its expression (Fig 3B, 3D). Expression of the neighboring gene *osbpl9*, encoding a lipid binding protein, was increased by 50% (Fig 3D), but *gas5^a173^* mutants were indistinguishable from wild type (Fig 3C), reached adulthood and were fertile.

**Figure 3:**
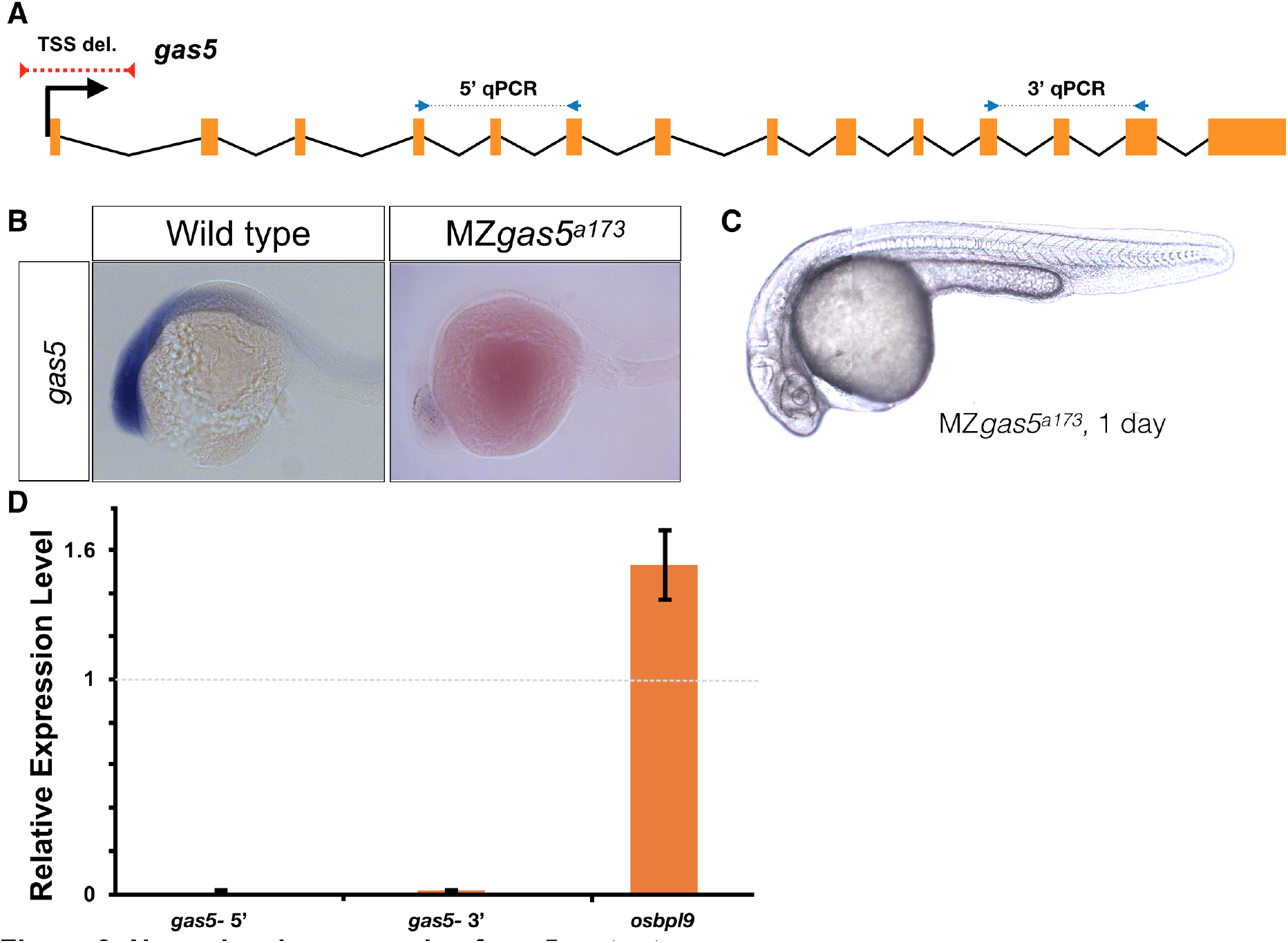
Normal embryogenesis of *gas5* mutants. A) Position of the TSS-deletion allele in gas5 is marked by dashed red line. Arrows flanking black dotted lines mark the primer binging sites for 5’-qPCR and 3’-qPCR products. B) Representative in situ hybridization images for gas5 in wild type (11/11) and homozygous TSS-deletion mutants (11/11). C) Maternal and Zygotic gas5 (MZgas5) mutant embryos at 1-dpf were indistinguishable from the wild-type embryos at the same developmental stage (not shown). D) Expression level of gas5 and osbpl9 measured by qRT-PCR. Tor3A, the other neighboring gene, was not expressed at the investigated time-point.

## Lnc-setd1ba

*Lnc-setd1ba* is the zebrafish orthologue of human LIMT ^43^ (LncRNA Inhibiting Metastasis), which has been implicated in basal-like breast cancers. It is expressed from a shared promoter region that also drives the expression of the histone methyltransferase *setd1ba* in opposite direction (Fig 4A). Evolutionary conservation in vertebrates and proximity to *setd1ba*, whose mouse homolog is essential for embryonic development^44,45^ prompted us to investigate the function of this lncRNA in zebrafish. We removed the gene of *lnc-setd1ba* downstream of its TSS (3137bp deletion) (*lnc-setd1ba*^a174^). In situ hybridization and qRT-PCR revealed absence of lncRNA expression (Fig 4C and 4E) and strong upregulation of *setd1ba* (Fig 4D and 4E) during cleavage stages and slight upregulation of *setd1ba* and the other neighboring gene *rhoF* at one-day post fertilization (1-dpf) (Fig 4E). Despite these changes, maternal-zygotic *lnc-setd1ba^a174^* mutants were indistinguishable from wild type (Fig 4B), reached adulthood and produced normal progeny.

**Figure 4:**
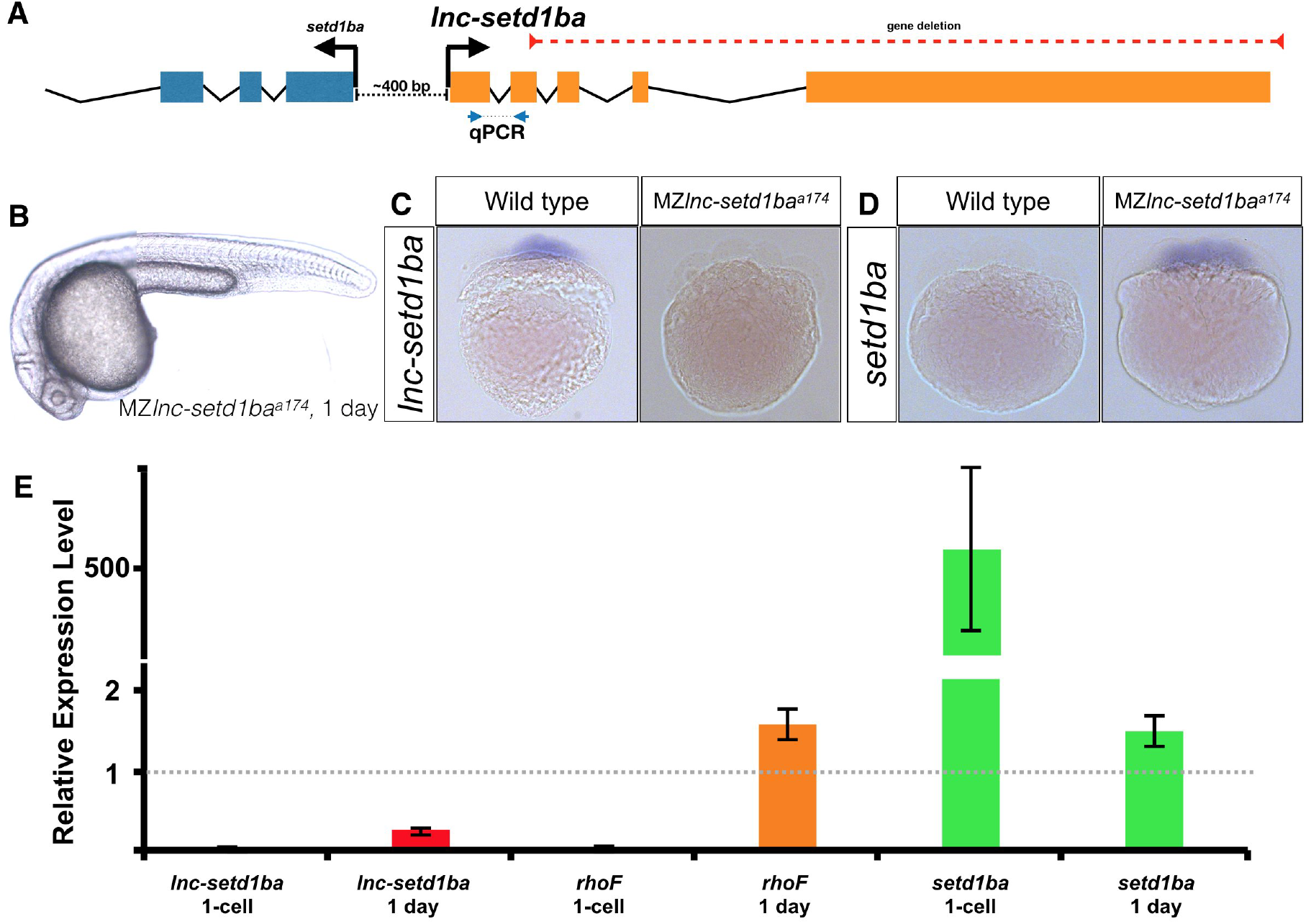
Normal embryogenesis of *lnc-setd1ba* mutants. A) The relative position of *lnc-setd1ba* and the protein-coding gene *setd1ba*. The gene deletion region is marked by dashed red line. Arrows flanking black dotted line mark the primer binging sites for qRT-PCR product. B) Maternal and zygotic *lnc-setd1ba* mutants were not different from wild-type embryos at 1-dpf. C) Representative images of in situ hybridization for *lnc-setd1ba* at 4-8 cell stage mutant (18/18) and wild-type (25/25) embryos. D) In situ hybridization for the protein-coding mRNA, *setd1ba* (9/11) in *lnc-setd1ba* mutants compared to the wild-type embryos (15/15). E) qRT-PCR at 1-cell stage and 1-dpf for the lncRNA and its neighboring genes *rhoF* and *setd1ba*.

## Squint

*Squint* encodes a Nodal ligand involved in mesendoderm specification^46,47^. The previously studied *squint* insertion mutant alleles (*squint^Hi975Tg 46^* and *squint^cz35 47^)* lead to delayed mesendoderm specification and partially penetrant cyclopia^48^. Morpholino and misexpression studies have suggested an additional, non-coding role for maternally provided s*quint*, wherein the *squint* 3′UTR mediates dorsal localization of *squint* mRNA, induces the expression of dorsal mesoderm genes, and is required for the development of dorsal structures^49,50^. This mode of activity assigns *squint* to the cncRNA family - RNAs with both protein-coding and non-coding roles^51^. To investigate the non-coding roles of *squint* mRNA we generated a deletion allele (*squint* ^a175^) that lacked most of the protein coding region and the 3’UTR, including the Dorsal Localization Element (DLE) implicated in maternal *squint* RNA localization^52^ (Fig 5A). In situ hybridization (Fig 5B) and qRT-PCR (Fig 5C) showed that the level of remaining *squint* transcript was greatly reduced (~90%). MZ*squint* ^a175^ embryos displayed partially penetrant cyclopia, similar to existing protein-disrupting *squint* alleles (Fig 5D)^46,47,53^, but the defects proposed to be caused by interference with *squint* non-coding activity^49^ were not detected.

**Figure 5:**
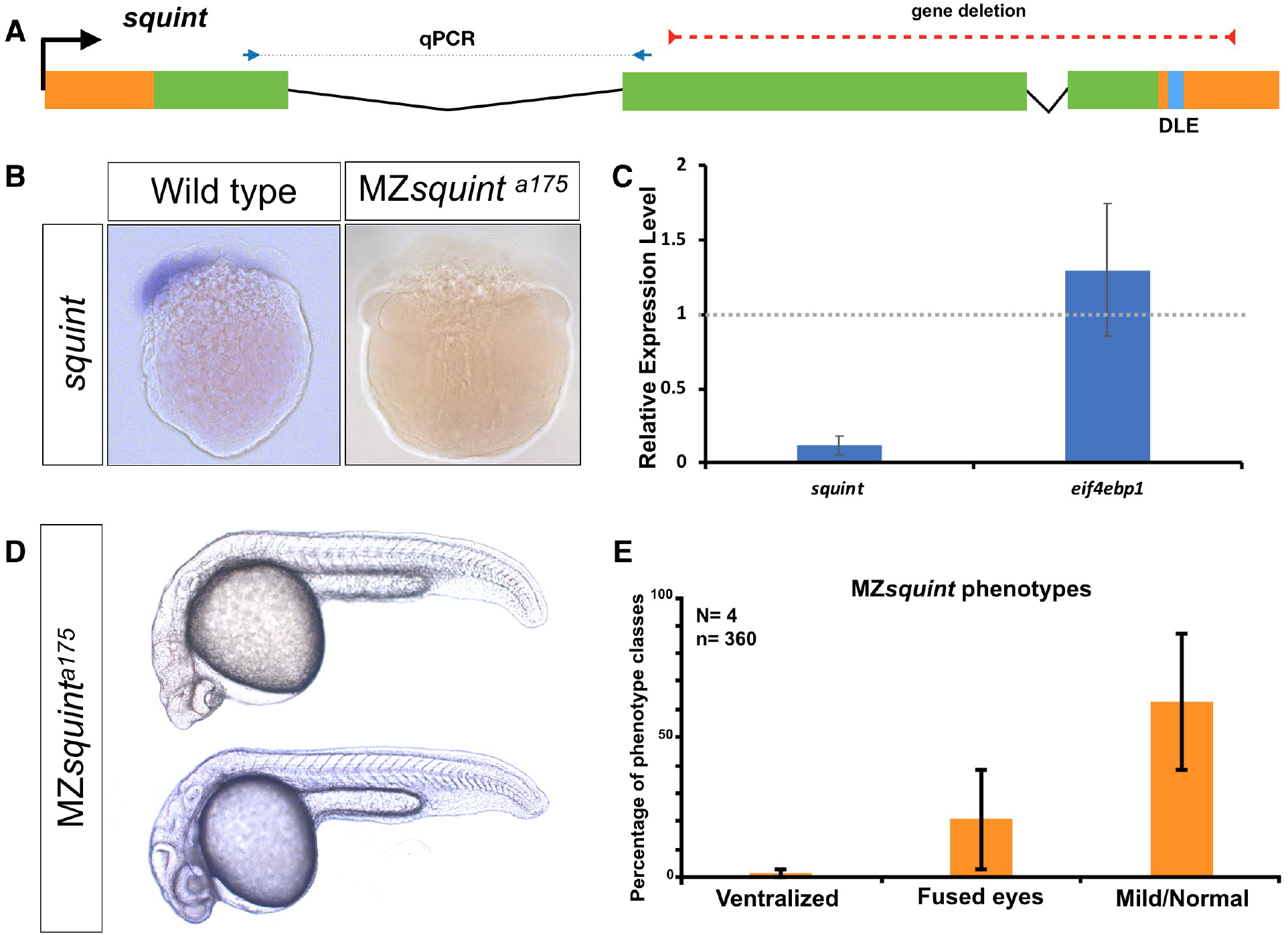
No non-coding function for *squint*. A) The position of untranslated regions (brown), coding region (green), putative Dorsal Localization Element-DLE (blue) and the gene deletion (red dashed line) in the *squint* genomic locus. Arrows flanking black dotted line mark the primer binging sites for qRT-PCR product. B) In situ hybridization for *squint* at 8-cell stage on wild-type (18/20) and MZ*squint^a175^*(17/17) embryos. C) MZ*squint^a175^* embryonic phenotype (N=4 independent crosses, n=360 embryos). D) Two representative MZ*squint^a175^* embryos. E) qRT-PCR for *squint* and *eif4ebp1* on wild-type and MZ*squint^a175^* embryos.

To further test whether *squint* mRNA might have non-coding roles, we injected wild-type and MZ*squint* ^a175^ embryos with either control RNA, full-length *squint* mRNA, a non-coding version of *squint* mRNA, or the putative transcript produced in *squint* ^a175^. We found that in contrast to wild-type *squint* mRNA, control RNA, non-protein coding *squint* RNA or *squint* ^a175^ RNA did not cause any phenotypes and did not rescue MZ*squint* ^a175^ mutants. These results indicate that *squint* does not have the previously proposed non-coding functions and is not a member of the cncRNA family.

## Transcript-independent phenotype at *lnc-phox2bb* locus

*Lnc-phox2bb* neighbors *phox2bb* and *smtnl1. Phox2bb* is a transcription factor implicated in the development of the sympathetic nervous system^54,55,56^, while *smtnl1* has been implicated in smooth muscle contraction ^57^. Whole-gene deletion of *lnc-phox2bb (lnc-phox2bb^a177^)* (Fig 6A) led to jaw deformation and failure to inflate the swim-bladder (Fig 6B), and no homozygous mutant fish survived to adulthood. Like the whole-gene deletion allele, the TSS-deletion allele (*lnc-phox2bb^a176^*) lacked *lnc-phox2bb* RNA (Fig S5 and Fig 6E), but in contrast to the whole-gene deletion mutants, TSS-deletion mutants developed normally and gave rise to fertile adults. To determine the cause of this difference, we analyzed the expression level and pattern of neighboring genes. We found that the anterior expression domain of *phox2bb* in the hindbrain was absent in the whole-gene deletion allele (Fig 6D). This finding is consistent with the observation that the deleted region contains enhancer elements for *phox2bb*^58^, conserved nongenic elements (CNEs)^59^, and histone marks related to enhancer regions (H3K4me1 and H3K27Ac)^60^. We also found that the expression level of *smtnl1* increased in gene deletion mutants relative to the TSS-deletion mutant and wild type (Fig 6E). These results indicate that *lnc-phox2bb* RNA is not required for normal development but that the *lnc-phox2bb* overlaps with regulatory elements required for proper expression of *phox2bb* and *smtnl1* (Fig 6E).

**Figure 6:**
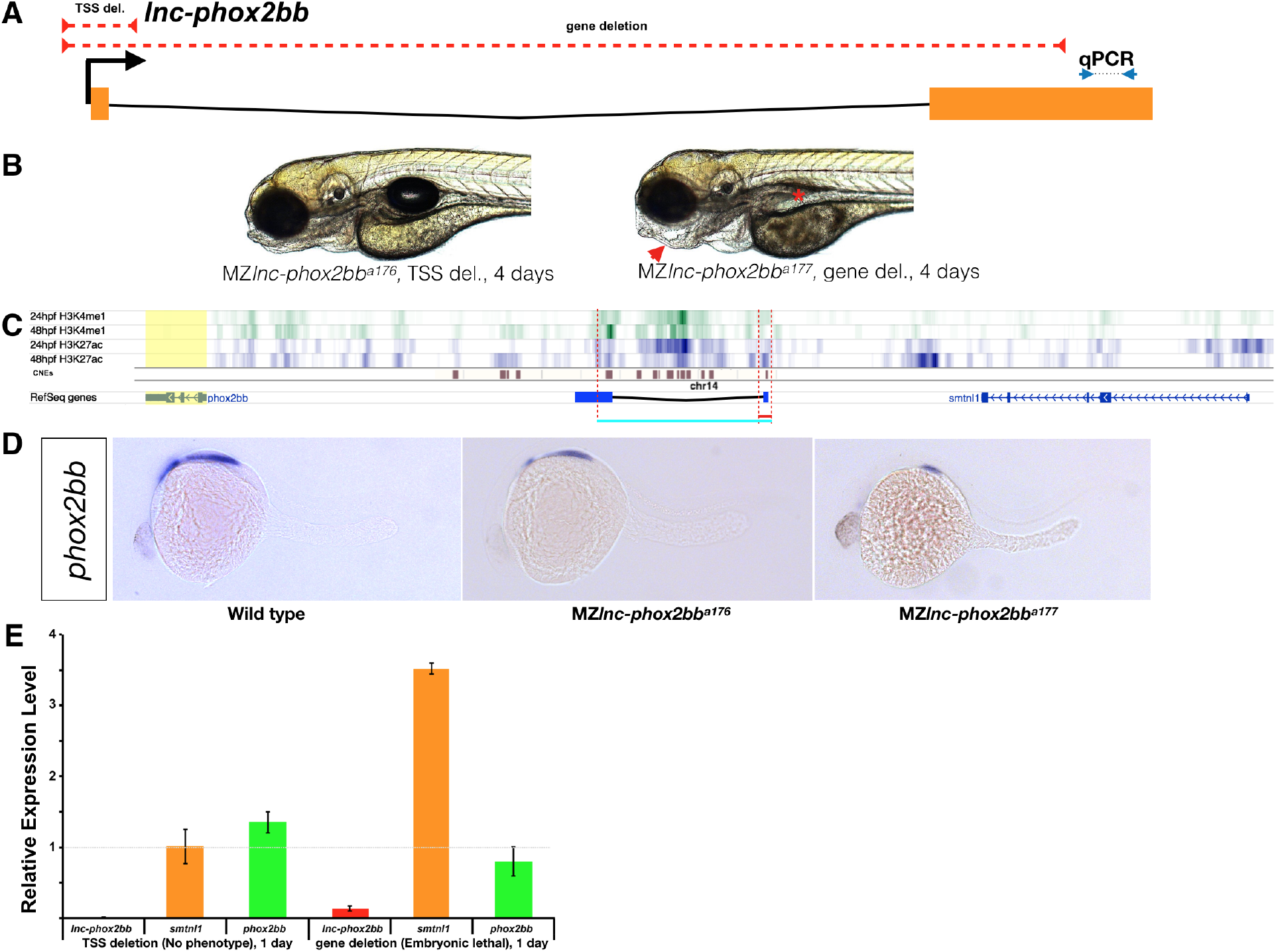
Requirement for *lnc-phox2bb* genomic elements but not RNA. A) The red dashed lines depict the respective positions of the *lnc-phox2bb* TSS and gene deletion. Arrows flanking black dotted line mark the primer binging sites for qRT-PCR product. B) Homozygous gene deletion mutants but not the TSS-deletion mutants show embryonic defects in jaw formation (arrow head) and swim bladder inflation (asterisk) by 4-dpf. C) Histone marks (H3K4me1 and H3K27ac) associated with enhancer activity^60^ and conserved non-genic elements CNEs^59^ overlap with gene deletion. D) *phox2bb* expression pattern in the TSS and gene deletions. E) qRT-PCR analysis on MZ TSS-deletion and gene deletion mutants.

In summary, our systematic mutant studies indicate that none of the 25 lncRNAs analyzed here are essential for embryogenesis, viability or fertility, including the prominent lncRNAs *cyrano*, *gas5*, and *lnc-setd1ba.* Additionally, they refute the proposed non-coding function of *squint* RNA. This mutant collection can now be analyzed for potentially context specific, redundant or subtle functions, but extrapolation suggests that most individual zebrafish lncRNAs are not required for embryogenesis, viability or fertility.

## Materials and Methods

### Animal care

TL/AB zebrafish (Danio rerio) were used as wild-type fish in this study. Fish were maintained on daily 14hr (light): 10hr (dark) cycle at 28°C. All animal work was performed at the facilities of Harvard University, Faculty of Arts & Sciences (HU/FAS). This study was approved by the Harvard University/Faculty of Arts & Sciences Standing Committee on the Use of Animals in Research & Teaching (IACUC; Protocol #25-08)

### Cas9 mediated mutagenesis

Guide RNAs (gRNAs) were designed using CHOPCHOP^61^ and synthesized in pool for each candidate as previously described^62^. (See supplementary file 1 for the gRNA sequences). gRNAs were combined with Cas9 protein (50 μM) and co-injected (~1 nL) into the one-cell stage TL/AB wild-type embryos. Genomic DNA from 10 injected and 10 un-injected siblings was extracted^63^ and screened for the difference in amplified band pattern from the targeted region (See supplementary file 1 for the genotyping primer sequences). The rest of injected embryos were raised to adulthood, crossed to wild-type fish and screened for passing the mutant allele to the next generation. Founder fish with desirable mutations were selected and confirmed by Sanger sequencing of the amplified mutant allele. Heterozygous mutants were crossed together to generate homozygous mutants. At least 15 adult homozygous mutant pairs per allele were crossed to test fertility of mutants and to generate maternal and zygotic mutants (MZ) devoid of maternally and zygotic lncRNA activity.

### Phenotype scoring procedure

Visual assessment of live embryos and larvae performed^30,31^ to identify mutant phenotypes, ranging from gastrulation movements and axis formation to the formation of brain, spinal cord, floor plate, notochord, somites, eyes, ears, heart, blood, pigmentation, vessels, kidney, pharyngeal arches, head skeleton, liver, and gut. At day five formation of swim bladder and overall appearance of the embryos were checked again (at any stage 60-100 embryos were scored). Sixty to hundred fish from heterozygous mutant crosses were grown to adulthood and genotyped to identify the viability of adult homozygous fish. Validated homozygous mutant fish were further crossed together to test for potential fertility phenotypes or putative maternal functions of candidate lncRNAs.

### Antisense RNA synthesis and in situ hybridization

Antisense probes for in situ hybridization were transcribed using the DIG RNA labeling kit (Roche). All RNAs were purified using EZNA Total RNA kits (Omega Biotek). Embryos were fixed in 4% formaldehyde overnight at 4°C (embryos younger than 50% epiboly fixed for 2 days). In situ hybridizations were performed according to standard protocols ^64^. NBT/BCIP/Alkaline phosphatase-stained embryos were dehydrated in methanol and imaged in benzyl benzoate:benzyl alcohol (BBBA) using a Zeiss Axio Imager.Z1 microscope.

### qRT-PCR

Total RNA was isolated from 3 individuals or sets of 10-20 embryos per condition using EZNA Total RNA kits (Omega Biotek). cDNA was generated using iScript cDNA Synthesis kit (Bio-Rad). qPCR was conducted using iTaq Universal SYBR Green Supermix (Bio-Rad) on a CFX96 (Bio-Rad). Gene expression levels were calculated relative to a reference gene, *ef1a*. The mean and standard deviation was plotted for each condition. Three technical replicates were used per condition. The qPCR primer sequences are listed in supplementary file 1.

### Bright field Imaging

Embryos were anesthetized in Tricaine (Sigma) and mounted in 1% low melting temperature agarose (Sigma) with Tricaine, then imaged using a Zeiss SteREO Discovery.V12 microscope or Zeiss Axio Imager.Z1 microscope. Images were processed in FIJI/ImageJ ^65^. Brightness, contrast and color balance was applied uniformly to images.

### Sense RNA synthesis and injection

The sequences for the wild-type *squint* mRNA, non-protein coding *squint* transcript (One Adenine base was added after 8 in-frame ATG codons, and the 3’UTR sequence kept unchanged) and the *squint^a175^* transcript were synthesized as gBlocks (IDT) containing 5’ XhoI cut site and 3’ NotI site. Fragments were digested and inserted the pCS2 plasmid. Positive colonies were selected, and sanger sequenced to assure the accuracy of the gene synthesis process. Sequences of the constructs are provided in supplementary file 1. mRNA was in vitro transcribed by mMessage mMachine (Ambion) and purified by EZNA Total RNA kits (Omega Biotek). H2B-GFP was used as control mRNA. Each injection mix contained 30ng/ul of squint or control mRNA). 1nl of mRNA mix was injected into the yolk of one-cell stage embryos.

## Acknowledgments

We thank current and former members of the Schier laboratory, particularly Andrea Pauli and Guo-Liang Chew for their helpful suggestions and support during the early phases of this project, Jeffrey Farrell, Nathan Lord and Maxwell Shafer for their critical comments on the manuscript, and the Harvard zebrafish facility staff for technical support. This work was supported by Leopoldina postdoctoral fellowship LPDS2014-01 to M.G. and NIH grant R01HD076708 to A.F.S.

## Author Contributions

M.G and A.F.S. designed the study, interpreted the data and wrote the manuscript. M.G. generated all data with support from K.B. and L.P.

---

**Figure S1:**
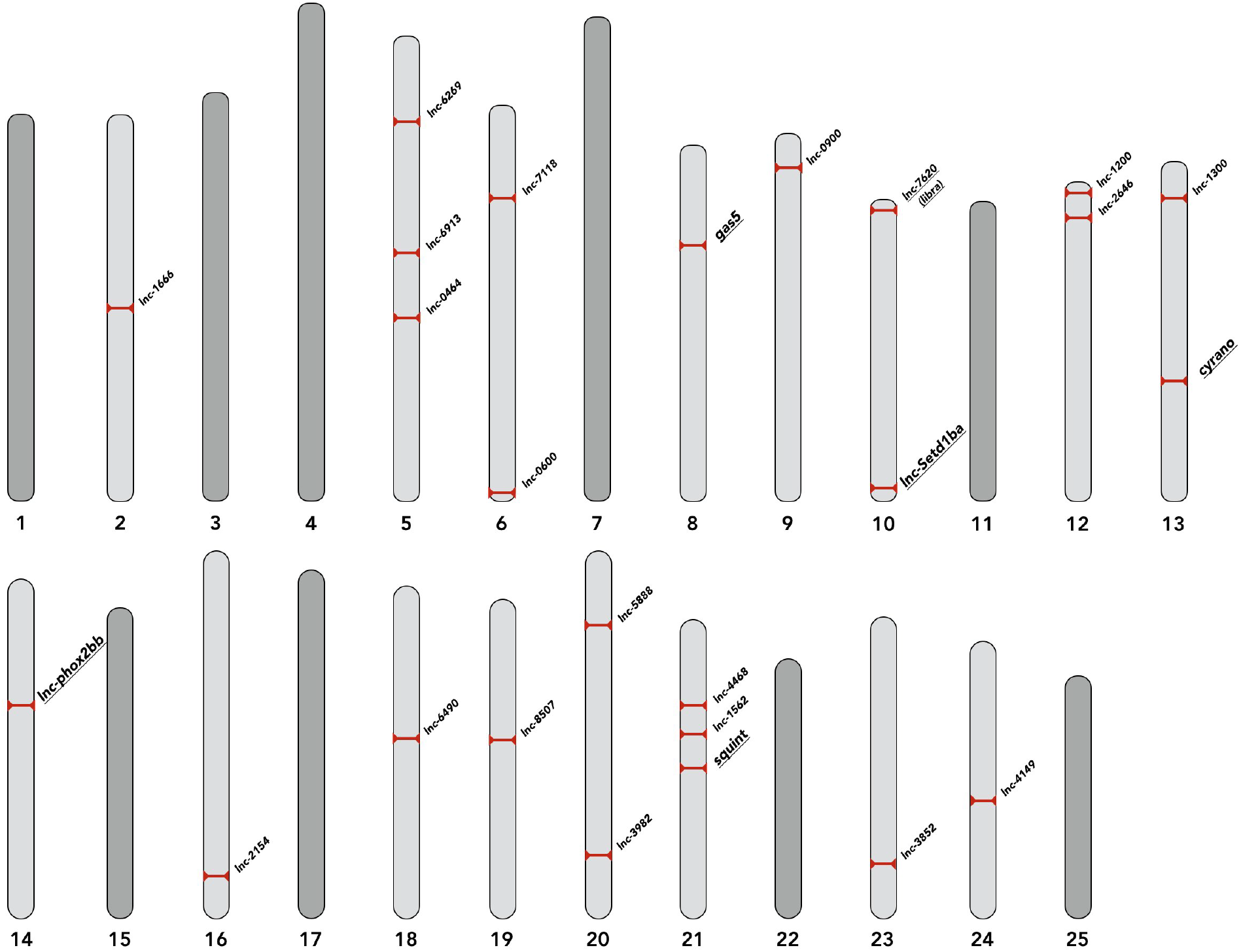
Genomic location of selected lncRNAs. The chromosomal positions of selected lncRNAs are depicted. lncRNAs discussed in the text are underlined.

**Figure S2:**
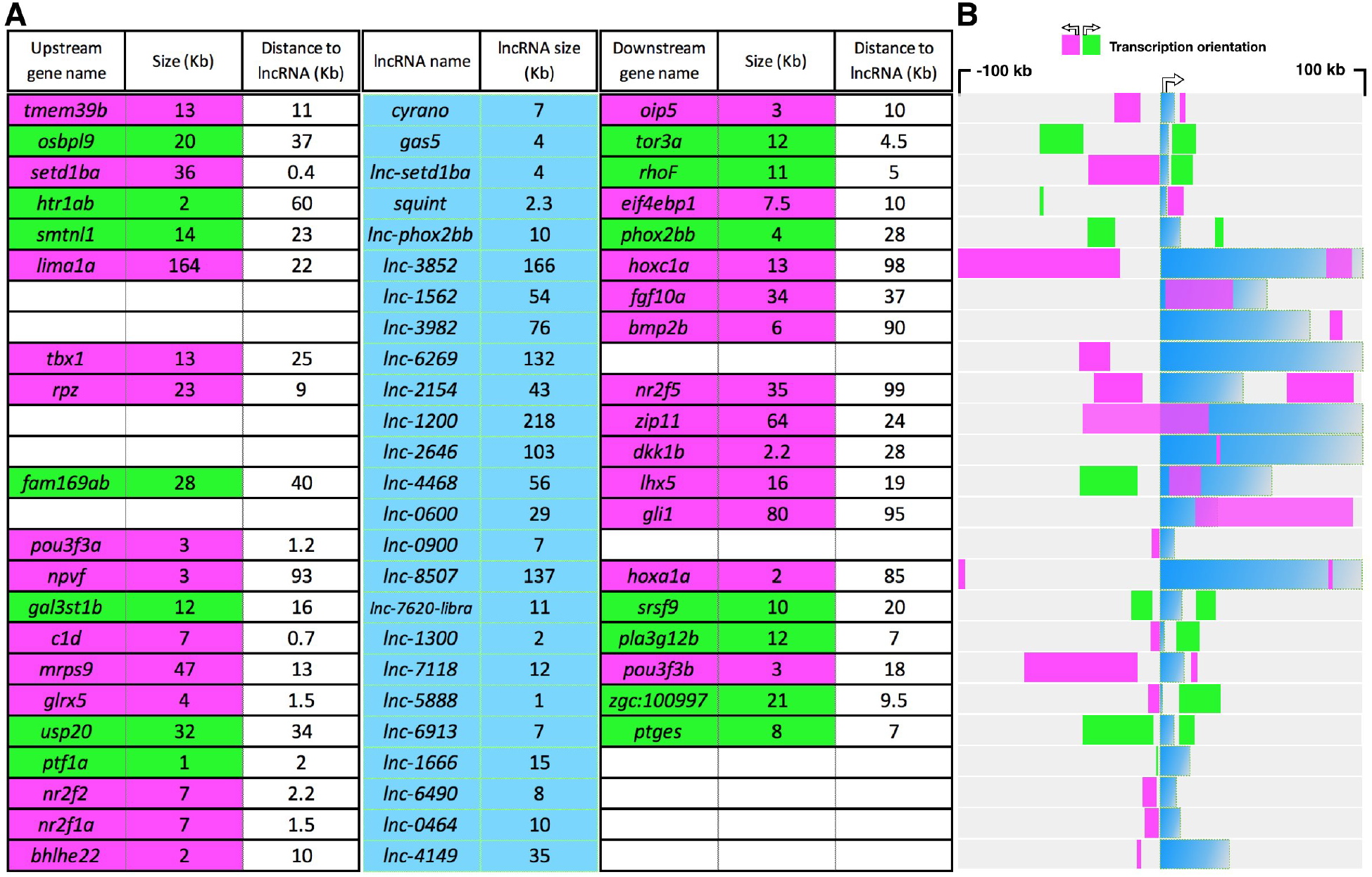
Size, relative distance and orientation of selected lncRNAs and their neighboring genes. A) lncRNA names and sizes are shown in the middle section (blue columns). The distance, size and transcriptional orientation of the neighboring genes, in a 200kb window centered on lncRNA’s TSS are shown on the left (upstream neighbor) and on the right (downstream neighbor). The transcription orientation is represented by green (in the same direction as lncRNA) and magenta (in the opposite direction of lncRNA). B) Visual representation of data in A. All sizes and distances are in Kb.

**Figure S3:**
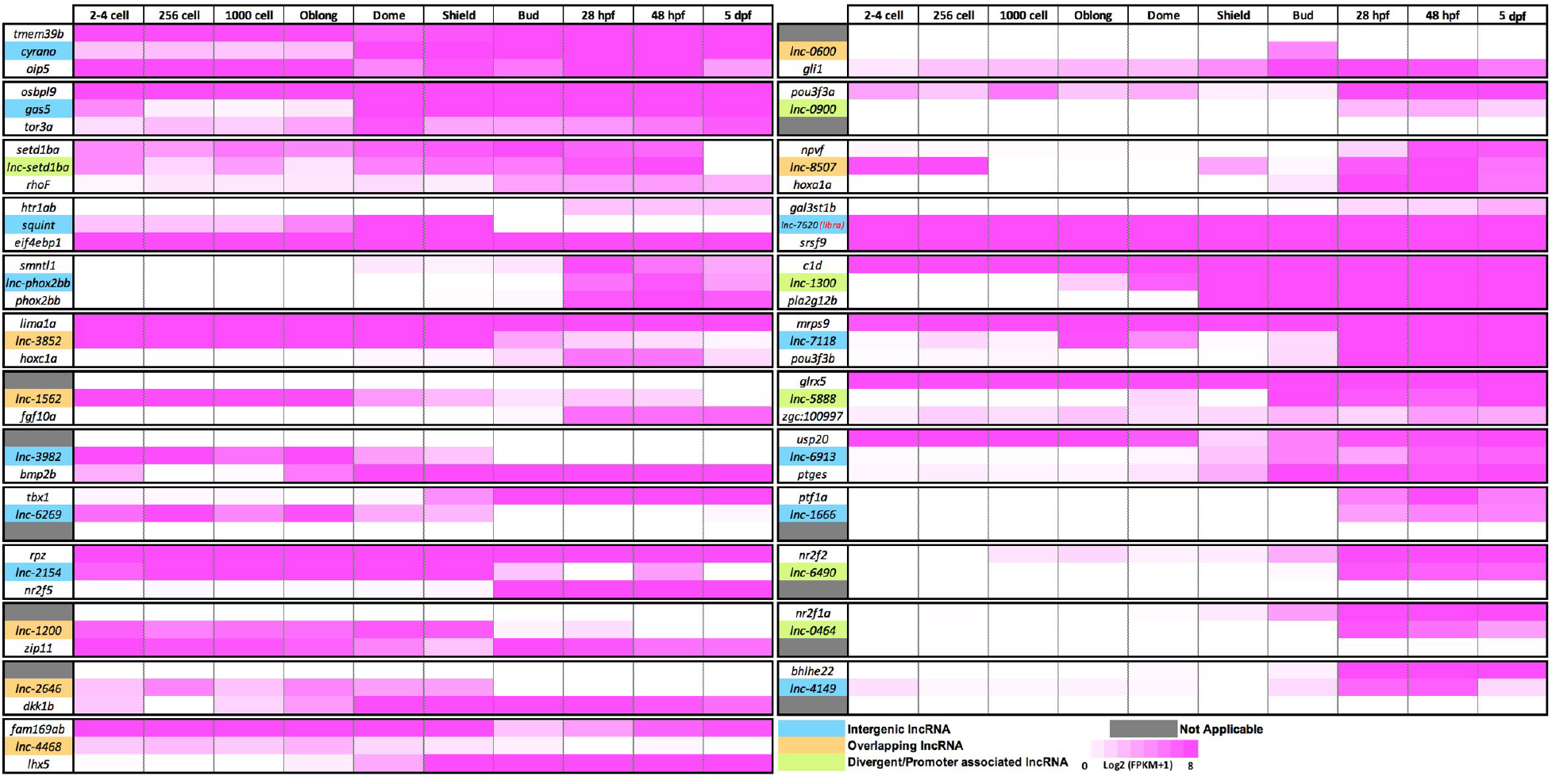
Expression levels of selected lncRNAs and their neighboring protein-coding genes. LncRNAs are color coded as blue (Intergenic), brown (Overlapping) and green (Divergent/Promoter associated) (see Fig S2B). For each lncRNA and its upstream (top) and downstream (bottom) neighbor, the expression levels at 10 early-developmental stages are shown^25^. The scale is log2 (FPKM+1) value, represented as gradient between 0 (white) and 8 (magenta).

**Figure S4:**
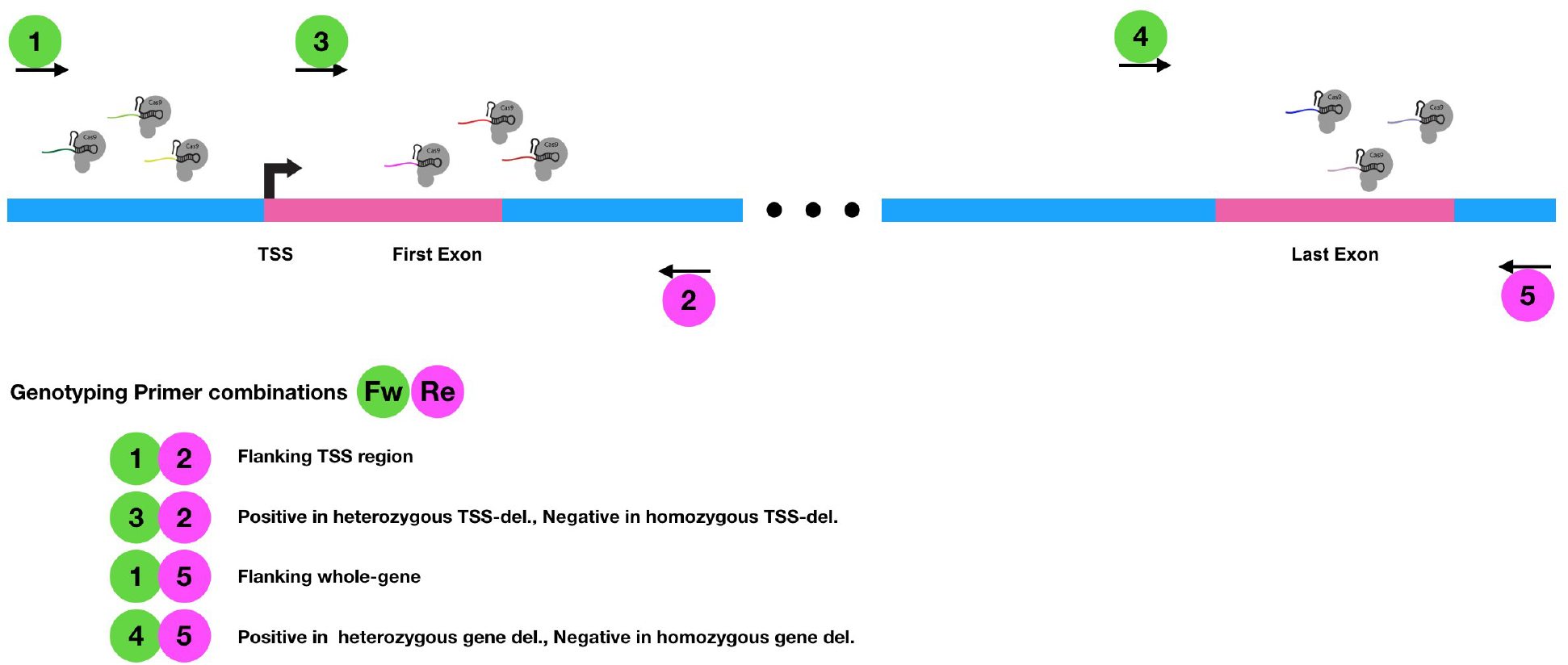
Cas9-mediated deletion approach for generating lncRNA knockouts. 6 gRNAs (three at either side of the TSS) were used to remove TSS. 9 guide RNAs (the first 6 plus three additional gRNAs around the Transcriptional Termination Site, TTS) were used to generate the gene deletions. Relative positions of genotyping primers are indicated by numbered circles.

**Figure S5:**
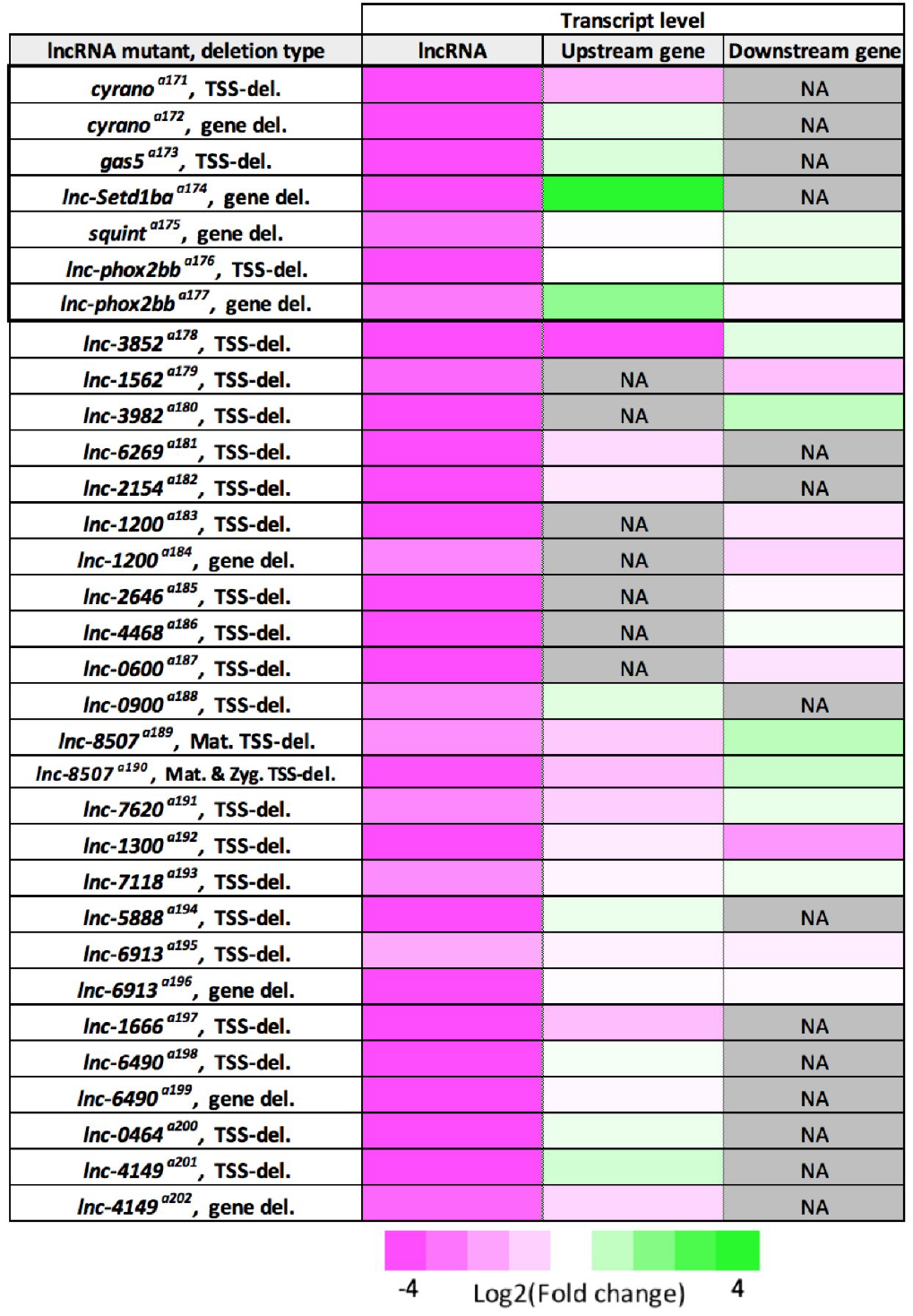
Summary of qRT-PCR analysis for lncRNA and their neighboring genes. Visual representation of the expression level changes for each lncRNA and its neighboring genes in homozygous deletion mutants. Three biological replicates for homozygous mutant and wild-type samples. Log2 of fold change between −4 (magenta) and 4 (green) is shown.

**Figure S6:**
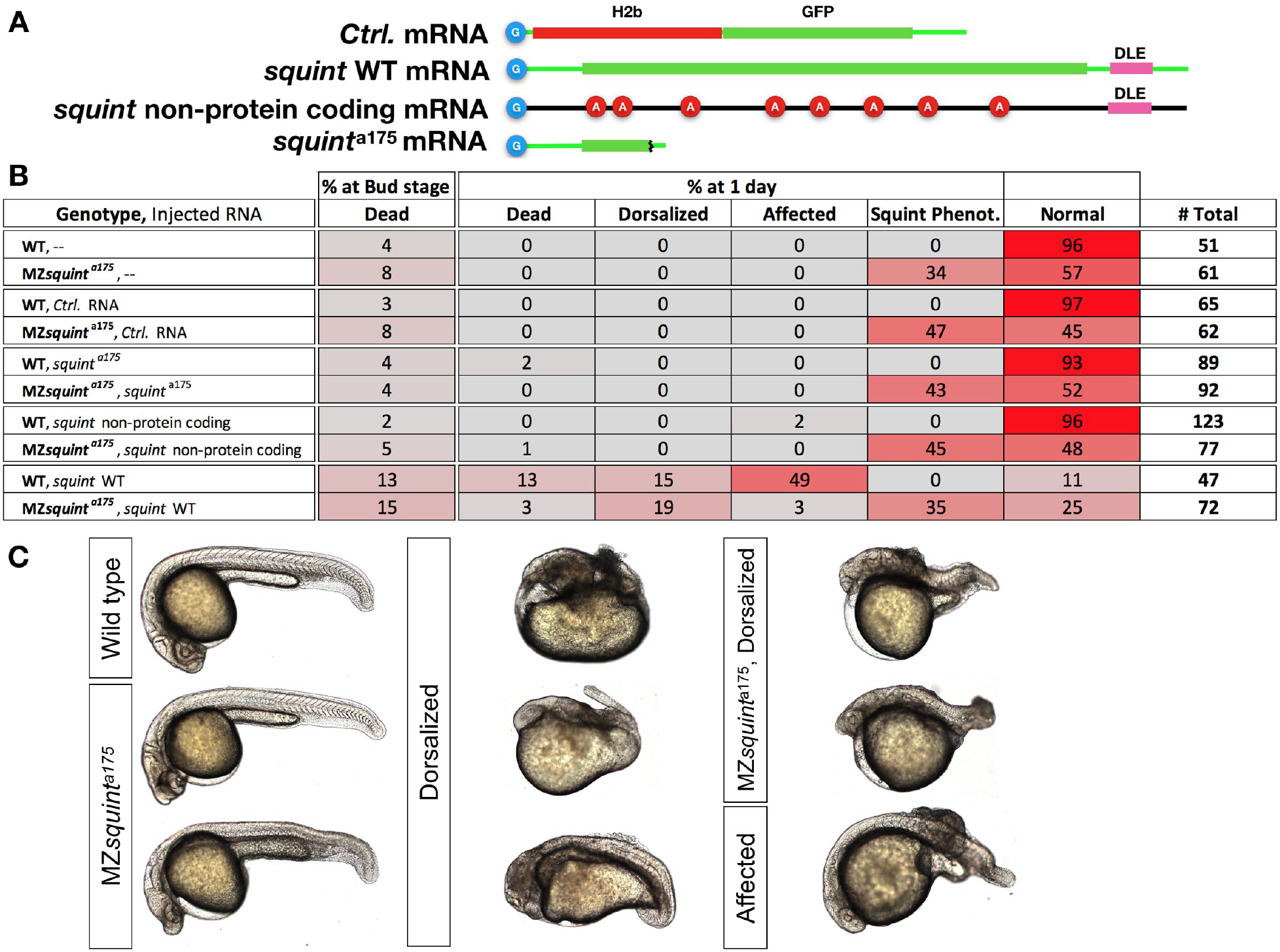
Dorsalization induced by Overexpression of *squint* mRNA but not its non-protein coding version. A) Schematic representation of injected mRNAs. Cap-analog is indicated by in blue circles at the beginning of each mRNA. *squint* non-protein coding mRNA was generated by adding 8 Adenine-nucleotides (red circles) after in-frame ATG codons. B) Table shows scoring outcome of observed phenotypes in embryos injected with 30pg of each indicated mRNA. C) Representative embryos showing typical wild-type, *squint* mutant or dorsalized morphology. Ambiguous phenotypes were scored as “Affected”.

**Supplementary File 1:**

This compressed folder contains three Excel files for the sequences of gRNAs, genotyping and qRT-PCR primers (for lncRNAs and their neighboring genes) and also the annotated sequence files (.ape) for each lncRNA and their deleted segments.

## References

1 St Laurent, G., Wahlestedt, C. & Kapranov, P. The Landscape of long noncoding RNA classification. Trends in genetics : TIG 31, 239–251, doi:10.1016/j.tig.2015.03.007 (2015).

2 Rinn, J. L. & Chang, H. Y. Genome regulation by long noncoding RNAs. Annual review of biochemistry 81, 145–166, doi:10.1146/annurev-biochem-051410- 092902 (2012).

3 Nagalakshmi, U. et al. The transcriptional landscape of the yeast genome defined by RNA sequencing. Science (New York, N.Y.) 320, 1344–1349, doi:10.1126/science.1158441 (2008).

4 Carninci, P. et al. The transcriptional landscape of the mammalian genome. Science (New York, N.Y.) 309, 1559–1563, doi:10.1126/science.1112014 (2005).

5 Kapranov, P. et al. RNA maps reveal new RNA classes and a possible function for pervasive transcription. Science (New York, N.Y.) 316, 1484–1488, doi:10.1126/science.1138341 (2007).

6 Brockdorff, N. et al. Conservation of position and exclusive expression of mouse Xist from the inactive X chromosome. Nature 351, 329–331, doi:10.1038/351329a0 (1991).

7 Marahrens, Y., Panning, B., Dausman, J., Strauss, W. & Jaenisch, R. Xist-deficient mice are defective in dosage compensation but not spermatogenesis. Genes & development 11, 156–166 (1997).

8 Lee, J. T., Davidow, L. S. & Warshawsky, D. Tsix, a gene antisense to Xist at the X-inactivation centre. Nature genetics 21, 400–404, doi:10.1038/7734 (1999).

9 Johnston, C. M., Newall, A. E., Brockdorff, N. & Nesterova, T. B. Enox, a novel gene that maps 10 kb upstream of Xist and partially escapes X inactivation. Genomics 80, 236–244 (2002).

10 Wutz, A. et al. Imprinted expression of the Igf2r gene depends on an intronic CpG island. Nature 389, 745–749, doi:10.1038/39631 (1997).

11 Latos, P. A. et al. Airn transcriptional overlap, but not its lncRNA products, induces imprinted Igf2r silencing. Science (New York, N.Y.) 338, 1469–1472, doi:10.1126/science.1228110 (2012).

12 Miyoshi, N. et al. Identification of an imprinted gene, Meg3/Gtl2 and its human homologue MEG3, first mapped on mouse distal chromosome 12 and human chromosome 14q. Genes to cells : devoted to molecular & cellular mechanisms 5, 211–220 (2000).

13 Kobayashi, S. et al. Mouse Peg9/Dlk1 and human PEG9/DLK1 are paternally expressed imprinted genes closely located to the maternally expressed imprinted genes: mouse Meg3/Gtl2 and human MEG3. Genes to cells : devoted to molecular & cellular mechanisms 5, 1029–1037 (2000).

14 Bartolomei, M. S., Zemel, S. & Tilghman, S. M. Parental imprinting of the mouse H19 gene. Nature 351, 153–155, doi:10.1038/351153a0 (1991).

15 Feil, R., Walter, J., Allen, N. D. & Reik, W. Developmental control of allelic methylation in the imprinted mouse Igf2 and H19 genes. Development (Cambridge, England) 120, 2933–2943 (1994).

16 Sauvageau, M. et al. Multiple knockout mouse models reveal lincRNAs are required for life and brain development. eLife 2, e01749, doi:10.7554/eLife.01749 (2013).

17 Anderson, K. M. et al. Transcription of the non-coding RNA upperhand controls Hand2 expression and heart development. Nature 539, 433–436, doi:10.1038/nature20128 (2016).

18 Amandio, A. R., Necsulea, A., Joye, E., Mascrez, B. & Duboule, D. Hotair Is Dispensible for Mouse Development. PLoS genetics 12, e1006232, doi:10.1371/journal.pgen.1006232 (2016).

19 Ip, J. Y. et al. Gomafu lncRNA knockout mice exhibit mild hyperactivity with enhanced responsiveness to the psychostimulant methamphetamine. Scientific reports 6, 27204, doi:10.1038/srep27204 (2016).

20 Bell, C. C. et al. The Evx1/Evx1as gene locus regulates anterior-posterior patterning during gastrulation. Scientific reports 6, 26657, doi:10.1038/srep26657 (2016).

21 Han, X. et al. Mouse knockout models reveal largely dispensable but context-dependent functions of lncRNAs during development. J Mol Cell Biol, doi:10.1093/jmcb/mjy003 (2018).

22 Lin, N. et al. An evolutionarily conserved long noncoding RNA TUNA controls pluripotency and neural lineage commitment. Molecular cell 53, 1005–1019, doi:10.1016/j.molcel.2014.01.021 (2014).

23 Ulitsky, I., Shkumatava, A., Jan, C. H., Sive, H. & Bartel, D. P. Conserved function of lincRNAs in vertebrate embryonic development despite rapid sequence evolution. Cell 147, 1537–1550, doi:10.1016/j.cell.2011.11.055 (2011).

24 Kok, F. O. et al. Reverse genetic screening reveals poor correlation between morpholino-induced and mutant phenotypes in zebrafish. Developmental cell 32, 97–108, doi:10.1016/j.devcel.2014.11.018 (2015).

25 Pauli, A. et al. Systematic identification of long noncoding RNAs expressed during zebrafish embryogenesis. Genome research 22, 577–591, doi:10.1101/gr.133009.111 (2012).

26 Kaushik, K. et al. Dynamic expression of long non-coding RNAs (lncRNAs) in adult zebrafish. PloS one 8, e83616, doi:10.1371/journal.pone.0083616 (2013).

27 Dhiman, H., Kapoor, S., Sivadas, A., Sivasubbu, S. & Scaria, V. zflncRNApedia: A Comprehensive Online Resource for Zebrafish Long Non-Coding RNAs. PloS one 10, e0129997, doi:10.1371/journal.pone.0129997 (2015).

28 Chew, G. L. et al. Ribosome profiling reveals resemblance between long non-coding RNAs and 5′ leaders of coding RNAs. Development (Cambridge, England) 140, 2828–2834, doi:10.1242/dev.098343 (2013).

29 Engreitz, J. M. et al. Local regulation of gene expression by lncRNA promoters, transcription and splicing. Nature 539, 452–455, doi:10.1038/nature20149 (2016).

30 Driever, W. et al. A genetic screen for mutations affecting embryogenesis in zebrafish. Development (Cambridge, England) 123, 37–46 (1996).

31 Haffter, P. et al. The identification of genes with unique and essential functions in the development of the zebrafish, Danio rerio. Development (Cambridge, England) 123, 1–36 (1996).

32 Sarangdhar, M. A., Chaubey, D., Srikakulam, N. & Pillai, B. Parentally inherited long non-coding RNA Cyrano is involved in zebrafish neurodevelopment. Nucleic acids research, doi:10.1093/nar/gky628 (2018).

33 Kim, J. et al. LncRNA OIP5-AS1/cyrano sponges RNA-binding protein HuR. Nucleic acids research 44, 2378–2392, doi:10.1093/nar/gkw017 (2016).

34 Smith, K. N., Starmer, J., Miller, S. C., Sethupathy, P. & Magnuson, T. Long Noncoding RNA Moderates MicroRNA Activity to Maintain Self-Renewal in Embryonic Stem Cells. Stem cell reports 9, 108–121, doi:10.1016/j.stemcr.2017.05.005 (2017).

35 Kleaveland, B., Shi, C. Y., Stefano, J. & Bartel, D. P. A Network of Noncoding Regulatory RNAs Acts in the Mammalian Brain. Cell, doi:10.1016/j.cell.2018.05.022 (2018).

36 Rossi, A. et al. Genetic compensation induced by deleterious mutations but not gene knockdowns. Nature 524, 230–233, doi:10.1038/nature14580 (2015).

37 El-Brolosy, M. A. & Stainier, D. Y. R. Genetic compensation: A phenomenon in search of mechanisms. PLoS genetics 13, e1006780, doi:10.1371/journal.pgen.1006780 (2017).

38 Coccia, E. M. et al. Regulation and expression of a growth arrest-specific gene (gas5) during growth, differentiation, and development. Molecular and cellular biology 12, 3514–3521 (1992).

39 Higa-Nakamine, S. et al. Loss of ribosomal RNA modification causes developmental defects in zebrafish. Nucleic acids research 40, 391–398, doi:10.1093/nar/gkr700 (2012).

40 Ma, C. et al. The growth arrest-specific transcript 5 (GAS5): a pivotal tumor suppressor long noncoding RNA in human cancers. Tumour biology : the journal of the International Society for Oncodevelopmental Biology and Medicine 37, 1437–1444, doi:10.1007/s13277-015-4521-9 (2016).

41 Pickard, M. R. & Williams, G. T. Molecular and Cellular Mechanisms of Action of Tumour Suppressor GAS5 LncRNA. Genes (Basel) 6, 484–499, doi:10.3390/genes6030484 (2015).

42 Schneider, C., King, R. M. & Philipson, L. Genes specifically expressed at growth arrest of mammalian cells. Cell 54, 787–793 (1988).

43 Sas-Chen, A. et al. LIMT is a novel metastasis inhibiting lncRNA suppressed by EGF and downregulated in aggressive breast cancer. EMBO molecular medicine 8, 1052–1064, doi:10.15252/emmm.201606198 (2016).

44 Eymery, A., Liu, Z., Ozonov, E. A., Stadler, M. B. & Peters, A. H. The methyltransferase Setdb1 is essential for meiosis and mitosis in mouse oocytes and early embryos. Development (Cambridge, England) 143, 2767–2779 (2016).

45 Kim, J. et al. Maternal Setdb1 is required for meiotic progression and preimplantation development in mouse. PLoS genetics 12, e1005970 (2016).

46 Pei, W., Williams, P. H., Clark, M. D., Stemple, D. L. & Feldman, B. Environmental and genetic modifiers of squint penetrance during zebrafish embryogenesis. Developmental biology 308, 368–378, doi:10.1016/j.ydbio.2007.05.026 (2007).

47 Heisenberg, C. P. & Nusslein-Volhard, C. The function of silberblick in the positioning of the eye anlage in the zebrafish embryo. Developmental biology 184, 85–94, doi:10.1006/dbio.1997.8511 (1997).

48 Dougan, S. T., Warga, R. M., Kane, D. A., Schier, A. F. & Talbot, W. S. The role of the zebrafish nodal-related genes squint and cyclops in patterning of mesendoderm. Development (Cambridge, England) 130, 1837–1851 (2003).

49 Gore, A. V. et al. The zebrafish dorsal axis is apparent at the four-cell stage. Nature 438, 1030–1035, doi:10.1038/nature04184 (2005).

50 Lim, S. et al. Dorsal activity of maternal squint is mediated by a non-coding function of the RNA. Development (Cambridge, England) 139, 2903–2915, doi:10.1242/dev.077081 (2012).

51 Sampath, K. & Ephrussi, A. CncRNAs: RNAs with both coding and non-coding roles in development. Development (Cambridge, England) 143, 1234–1241, doi:10.1242/dev.133298 (2016).

52 Gilligan, P. C. et al. Conservation defines functional motifs in the squint/nodal-related 1 RNA dorsal localization element. Nucleic acids research 39, 3340–3349, doi:10.1093/nar/gkq1185 (2011).

53 Golling, G. et al. Insertional mutagenesis in zebrafish rapidly identifies genes essential for early vertebrate development. Nature genetics 31, 135–140, doi:10.1038/ng896 (2002).

54 Pei, D. et al. Distinct neuroblastoma-associated alterations of PHOX2B impair sympathetic neuronal differentiation in zebrafish models. PLoS genetics 9, e1003533 (2013).

55 Moreira, T. S., Takakura, A. C., Czeisler, C. & Otero, J. J. Respiratory and autonomic dysfunction in congenital central hypoventilation syndrome. J Neurophysiol 116, 742–752, doi:10.1152/jn.00026.2016 (2016).

56 Tolbert, V. P., Coggins, G. E. & Maris, J. M. Genetic susceptibility to neuroblastoma. Curr Opin Genet Dev 42, 81–90, doi:10.1016/j.gde.2017.03.008 (2017).

57 Borman, M. A., Freed, T. A., Haystead, T. A. & Macdonald, J. A. The role of the calponin homology domain of smoothelin-like 1 (SMTNL1) in myosin phosphatase inhibition and smooth muscle contraction. Mol Cell Biochem 327, 93–100, doi:10.1007/s11010-009-0047-z (2009).

58 McGaughey, D. M. et al. Metrics of sequence constraint overlook regulatory sequences in an exhaustive analysis at phox2b. Genome research 18, 252–260, doi:10.1101/gr.6929408 (2008).

59 Hiller, M. et al. Computational methods to detect conserved non-genic elements in phylogenetically isolated genomes: application to zebrafish. Nucleic acids research 41, e151, doi:10.1093/nar/gkt557 (2013).

60 Bogdanovic, O. et al. Dynamics of enhancer chromatin signatures mark the transition from pluripotency to cell specification during embryogenesis. Genome research 22, 2043–2053, doi:10.1101/gr.134833.111 (2012).

61 Montague, T. G., Cruz, J. M., Gagnon, J. A., Church, G. M. & Valen, E. CHOPCHOP: a CRISPR/Cas9 and TALEN web tool for genome editing. Nucleic acids research 42, W401–407, doi:10.1093/nar/gku410 (2014).

62 Gagnon, J. A. et al. Efficient mutagenesis by Cas9 protein-mediated oligonucleotide insertion and large-scale assessment of single-guide RNAs. PloS one 9, e98186, doi:10.1371/journal.pone.0098186 (2014).

63 Meeker, N. D., Hutchinson, S. A., Ho, L. & Trede, N. S. Method for isolation of PCR-ready genomic DNA from zebrafish tissues. Biotechniques 43, 610, 612, 614 (2007).

64 Thisse, C. & Thisse, B. High-resolution in situ hybridization to whole-mount zebrafish embryos. Nat Protoc 3, 59–69, doi:10.1038/nprot.2007.514 (2008).

65 Schindelin, J. et al. Fiji: an open-source platform for biological-image analysis. Nat Methods 9, 676–682, doi:10.1038/nmeth.2019 (2012).

